# TMS timed to interictal epileptiform discharges

**DOI:** 10.64898/2026.02.17.706146

**Authors:** Matilda Makkonen, Olli-Pekka Kahilakoski, Miguel Menchaca, Ivan Zubarev, Oula Siljamo, Umair Hassan, Xiwei She, Wendy Qi, Tuomas P. Mutanen, Risto J. Ilmoniemi, Pantelis Lioumis, Fiona M. Baumer

## Abstract

Interictal epileptiform discharges (IEDs) are pathological hypersynchronous bursts of electrical brain activity that occur between seizures in patients with epilepsy. IEDs are caused by transient brain states that are difficult to predict, making them a challenging neurophysiological and technological case for brain-state-dependent stimulation. Administering stimulation at IED onset may provide insight into the epileptic network and optimize neurostimulation therapies. Here, we assessed the feasibility of IED-triggered transcranial magnetic stimulation (TMS) in two children with self-limited epilepsy with centrotemporal spikes (SeLECTS), a common pediatric epilepsy in which IEDs emerge from the motor cortex.

A convolutional neural network (CNN) was trained on the participants’ pre-recorded electroencephalography (EEG) data with IEDs annotated by an epileptologist. The CNN was integrated into an EEG-processing pipeline that classified EEG segments as “IED” or “non-IED” in real time. With this pipeline, TMS pulses were administered during IED or non-IED periods in an interleaved, randomized design. We stimulated both the motor cortex generating the IEDs and the contralateral motor cortex and tested the impact of IEDs on TMS-evoked potentials (TEPs).

Our study demonstrated that TMS can be timed to IEDs and that there is a site-specific increase in TEP amplitude when stimulating during IEDs. Out of the TMS pulses aimed at an IED, 39% and 19% were successfully delivered during an IED for the two participants, respectively. For future research, we propose ways to address the methodological challenges of IED-timed TMS, enabling brain-state-dependent TMS for epilepsy research and treatment.

## 1. Introduction

Interictal epileptiform discharges (IEDs) are brief (<200 ms) hypersynchronous bursts of electrical brain activity in patients with epilepsy, detectable with electroencephalography (EEG). IEDs and seizures emerge from a transient, unpredictable brain state associated with excitation-inhibition imbalance, often requiring prolonged inpatient EEG monitoring for accurate characterization. Thus, new technology to study IEDs and similar transient brain states would offer substantial clinical benefit.

Transcranial magnetic stimulation (TMS) is a non-invasive method for activating cortical neurons that can be paired with EEG (TMS–EEG) to measure TMS-evoked potentials (TEPs), time-locked positive and negative deflections. The main TEP component observed in children is a negative deflection 100 ms post-stimulus (N100) (Baumer et al., 2020; Bender et al., 2005). TEPs are influenced by local cortical excitability and reactivity, and their propagation reflects local and global brain dynamics (Bender et al., 2005) and can support epilepsy diagnosis (Kimiskidis et al., 2017). Repetitive TMS (rTMS) can decrease cortical excitability and hyperconnectivity, offering therapeutic potential in epilepsy (VanHaerents et al., 2020). Responses to both single and rTMS pulses vary depending on the instantaneous brain state (Bradley et al., 2022), motivating development of methods to target specific brain states.

IEDs make an interesting neurophysiological and technological case for brain-state-dependent TMS, enabling interrogation of epileptic networks and optimization of rTMS therapies. Algorithms to detect and stimulate IEDs have been implemented using invasive, intracranial EEG (Zelmann et al., 2020), but scalp EEG presents unique challenges due to lower signal-to-noise ratio and inferior spatial resolution. Non-invasive detection and stimulation approaches, however, (Allen et al., 2017) offer more widespread applicability for research and treatment.

We demonstrate the feasibility of IED-timed TMS measurements in two children with self-limited epilepsy with centrotemporal spikes (SeLECTS), a common focal pediatric epilepsy (Specchio et al., 2022). This syndrome is a compelling test case for IED-timed TMS, as patients have frequent IEDs from the motor region (Specchio et al., 2022), which is easily targeted with TMS. We present a novel preprocessing method for IED-timed TMS−EEG and explore the effect of IEDs on TEP size and propagation. We hypothesize that TEPs will propagate more during IED than non-IED states due to increased brain connectivity and excitability.

## 2. Methods

### 2.1. Participants

Two 11-year-old right-handed boys with SeLECTS were studied with Stanford University Institutional Review Board (no. 68956) approval. Written consent and assent were obtained from parents and participants, respectively. We refer to participants as P1 and P2. P1 was not taking daily antiseizure medications, while P2 was taking daily oxcarbazepine (dose 10 mg/kg/day; serum level 22). Both participants had frequent centrotemporal IEDs when awake, P1 with bilateral (but left-hemisphere predominant) and P2 with exclusively right-hemisphere IEDs.

### 2.2. IED-detection algorithm

An IED-detecting machine-learning model was trained separately for each participant. Training EEG data were recorded with a BrainVision actiCHamp Pulse amplifier and a 64-channel cap with slim active electrodes, sampled at 25 kHz. Duration of the training-data recording was 1134 s for P1 and 959 s for P2, from which 327 (P1) and 498 (P2) IEDs were annotated. We trained the linear-filter convolutional neural network (LF-CNN) algorithm (Zubarev et al., 2019) to classify 130-ms EEG segments as “IED” or “non-IED.” The trained model was integrated into real-time EEG-guided TMS software NeuroSimo (Kahilakoski et al., 2025), which classified the latest 130 ms of data every 20 ms. Details of training data, procedures, and algorithm are in the Supplementary Material.

### 2.3. TMS–EEG measurements and real-time setup

TMS−EEG measurements were performed at 10 am, three months (P1) and one week (P2) after training data acquisition. Four regions of interest (ROIs) were stimulated: left and right primary motor cortices (M1) were primary ROIs, whereas left and right occipital cortices were control ROIs as they are far from M1 and not thought to be directly involved in SeLECTS (Specchio et al., 2022). The Localite TMS neuronavigation system with an age-appropriate magnetic resonance image (MRI) template (Fonov et al., 2011) ensured consistent administration of stimulation.

M1 targets were the motor hotspots, *i.e*., cortical areas evoking the largest motor-evoked potentials (MEPs) for left and right *first dorsal interosseous* (FDI) muscles. We identified the resting motor threshold (rMT) of each hemisphere, defined as the minimum intensity evoking MEPs with peak-to-peak amplitude ≥50 μV in 5 of 10 pulses. For P1, rMTs were 80% (left) and 83% (right) of the maximum stimulator output (MSO). For P2, for whom no MEPs were elicited at rest, M1 targets were identified during mild hand contraction. Occipital ROIs were defined as equidistant from midline around the POz electrode. No phosphenes were elicited. Stimulation intensity for all sites was 120% rMT, up to 100% MSO. Thus, P1 received suprathreshold stimulation (120% rMT was 96−100% MSO), whereas P2 did not (true rMT exceeded 100% MSO).

Real-time EEG was recorded using the same cap and amplifier as the training data, sampled at 1 kHz. A TurboLink device streamed EEG from the amplifier to a laptop computer running NeuroSimo, which generated and transmitted trigger signals to a MagVenture MagPro X100 TMS device via a LabJack T4 USB peripheral. Stimulation was delivered in blocks of up to 99 pulses, totaling 925 (P1) and 693 (P2) pulses across all sites. Each block consisted of matched numbers of stimuli aimed at “IED” and “non-IED” states in a pre-defined, randomized order, aimed at the selected state based on the latest EEG classification. The minimum inter-stimulus interval (ISI) was 2 s for P1. For P2, the minimum ISI was shortened to 1 s before IED stimuli, and to 1.5 s after the previous IED detection for non-IED stimuli (rationale and data acquisition details in Supplementary Material).

### 2.4. Data processing

#### 2.4.1. IED annotation & detection algorithm evaluation

After the real-time TMS experiment, EEG was epoched around each TMS pulse (−1000 ms to +1500 ms for P1; −950 to +950 ms for P2, due to different ISIs). IEDs occurring within these epochs were annotated by a pediatric epileptologist (FMB). Algorithm performance was evaluated by comparing the classification output (IED or non-IED) against actual presence of IEDs from −50 ms to +20 ms around each TMS pulse. Epochs were grouped into four categories (Supplementary Figs. S1 and S2): true positive (classified as IED / IED present), false positive (classified as IED / no IED present), true negative (classified as non-IED / no IED present), or false negative (classified as non-IED, IED present). Clopper-Pearson confidence intervals (CI) were calculated for proportions of true positives out of all positives and true negatives out of all negatives. IED-detection algorithm success rate was compared with the theoretical probability of hitting IEDs with randomly timed pulses (Supplementary Material).

#### 2.4.2. Extraction of TMS-evoked potentials

##### 2.4.2.1. Epoch selection

We defined three conditions: IED−TEP (true positives); non-IED TEP (nTEP; true negatives); and IED-alone (IEDs occurring >1 s from TMS pulses and other IEDs). To create IED-alone epochs, we placed a marker after the maximum negativity of each IED, matched to where the TMS pulse fell within IED−TEP epochs (P1 at 20 ms; P2 at 7 ms post-pulse). For all conditions, we excluded epochs with IEDs between −500 to −50 ms or +20 to +300 ms of the TMS pulse to limit IED effects on the baseline or TEPs.

##### 2.4.2.2. Preprocessing

We developed a preprocessing pipeline for TEPs containing IEDs that distort the signal in standard pipelines (Fig. 1; Supplementary Material). For each participant, epochs from the three conditions were concatenated and preprocessed together to maintain comparable preprocessing-induced signal across conditions. Data from each stimulation site were preprocessed separately due to different stimulation-induced artifacts. Pipeline steps included (Fig. 1): removing and interpolating the TMS pulse artifact; removing low-frequency drifts with robust detrending (de Cheveigné & Arzounian, 2018); manually discarding artifactual epochs; matching epoch numbers across conditions (IED−TEP condition had fewest epochs); removing ocular artifacts with independent component analysis (ICA); suppressing noise with the source-estimate-utilizing noise-discarding (SOUND) algorithm (Mutanen et al., 2022); removing muscle artifacts with signal-space-projection–source-informed reconstruction (SSP–SIR) (Mutanen et al., 2022); and low-pass filtering.

**Figure 1.**
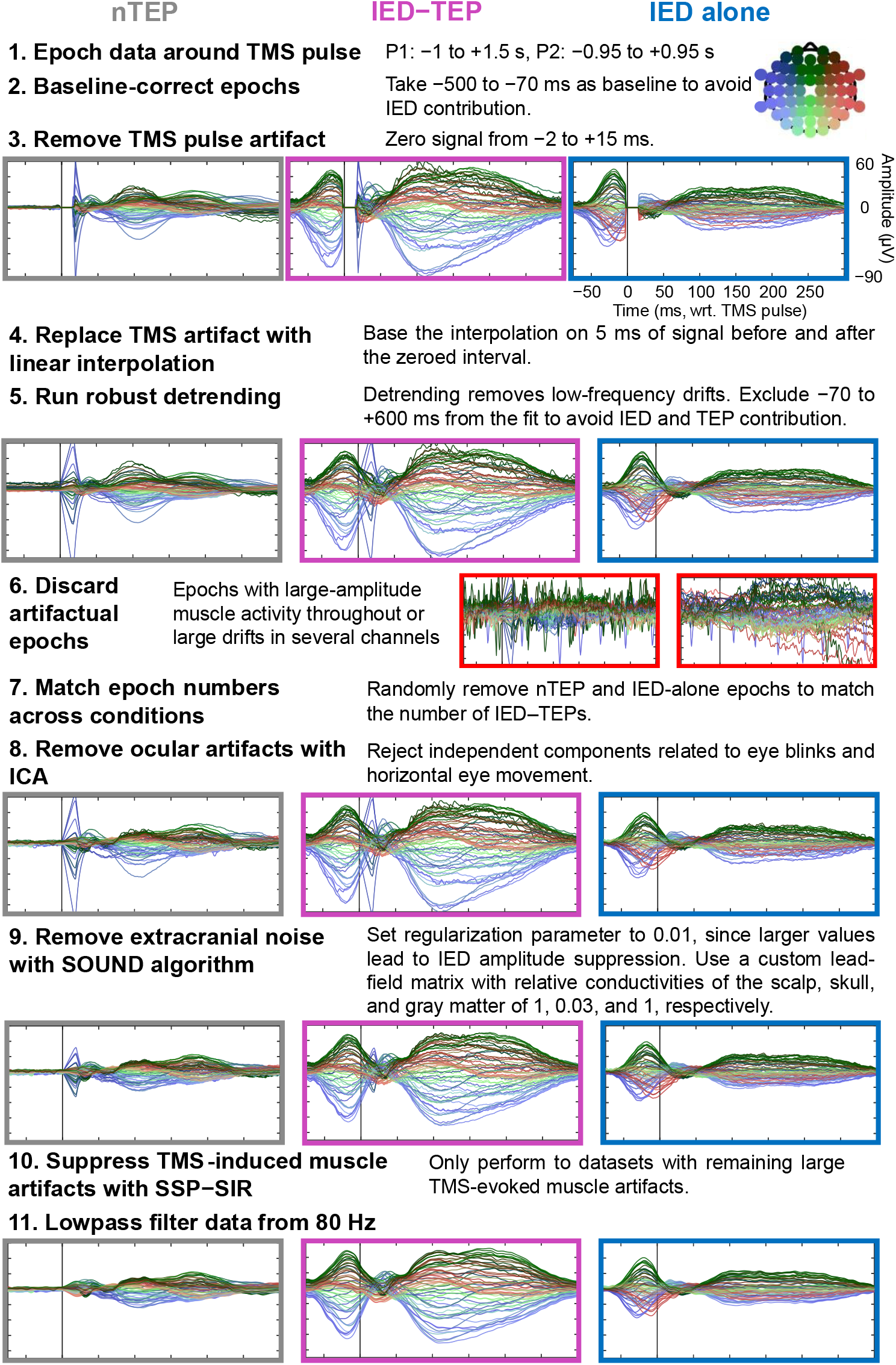
Preprocessing pipeline with examples of intermediate steps. Representative signals presented from left M1 stimulation of P1 are shown in the global average reference. Data from all three conditions, nTEP (grey), IED−TEP (pink), and IED-alone (blue), were preprocessed together. Step 8 forward averages 23 epochs per condition; for earlier steps, number of epochs averaged depends on condition.

The SOUND algorithm was employed with a custom lead-field matrix built from the MRI template used for neuronavigation. When computing lead fields, relative conductivities of the scalp, skull, and gray matter were set to 1, 0.03, and 1, accounting for the higher skull conductivity in children versus adults (McCann & Beltrachini, 2021).

#### 2.4.3. TEP analysis

For each condition, we calculated global mean field amplitudes (GMFA) from all electrodes and local mean field amplitudes (LMFA) for ROI electrodes (Figs. 2 and S3). From LMFA and GMFA, we calculated the area under the curve (AUC) for the late TEP (80–200 ms), corresponding to the most prominent component in pediatric TEPs, the N100 (Baumer et al., 2020; Bender et al., 2005). AUCs across conditions were compared with the Kruskal-Wallis (KW) non-parametric test with Dunn–Šidák post-hoc test to identify specific pairwise differences. Separate KWs were run for LMFA and GMFA for each stimulation site.

**Figure 2.**
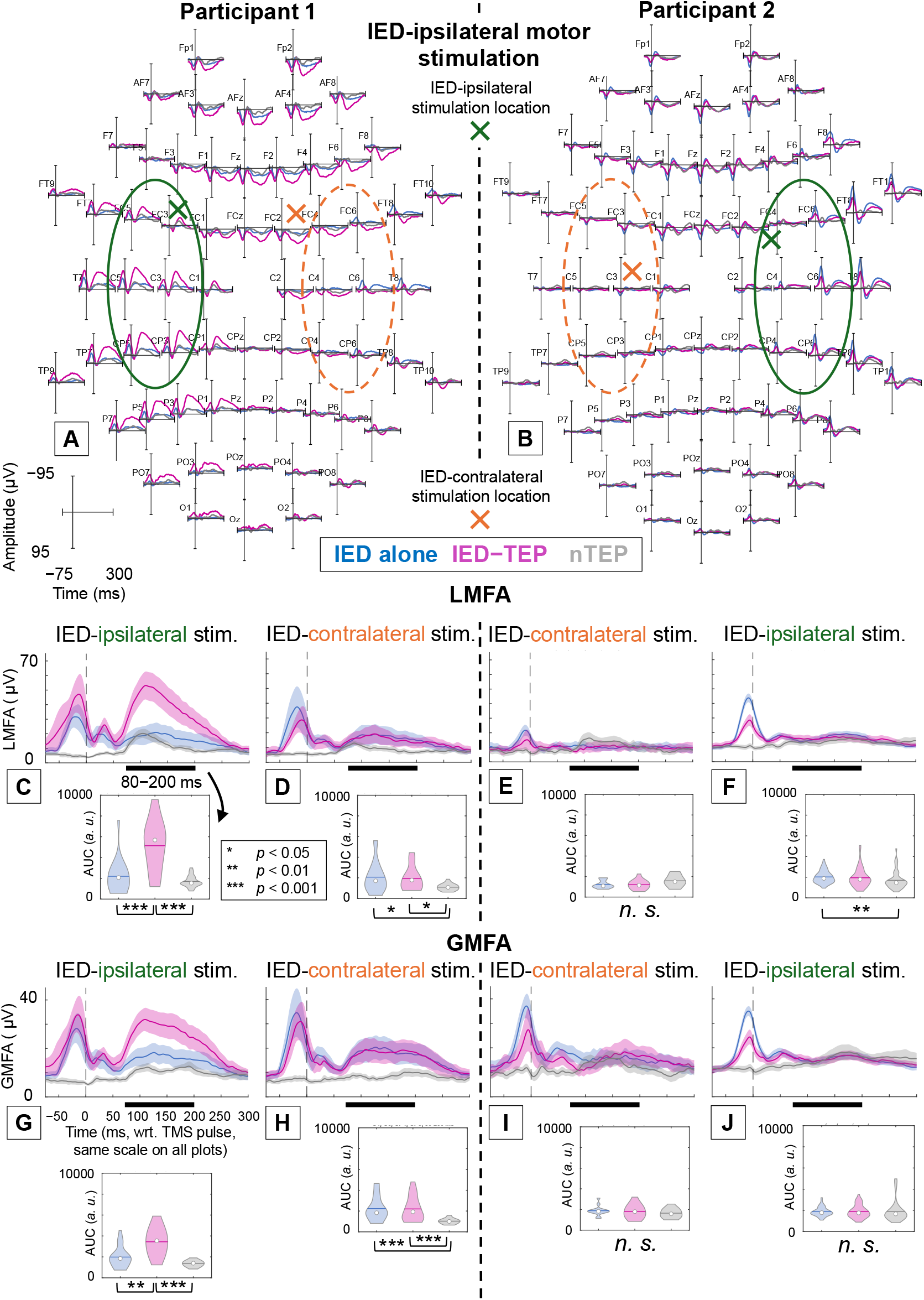
Panels A and B: Topographic plots of EEG signals in each channel in the global average reference for IEDs (blue), and TMS-evoked responses when successfully targeting an IED (pink) and a non-IED (nTEP, grey) for P1 (A) and P2 (B); stimulation applied to IED-generating (ipsilateral) motor cortex (A: left M1, B: right M1). Stimulation sites are presented as crosses with ROI electrodes circled. Green cross and circle represent ipsilateral stimulation and ROI, respectively, while orange cross and circle represent the contralateral counterparts. Panels C to F: Local mean field amplitudes (LMFA) of the six-electrode ROI. Panels G to J: Global mean field amplitudes (GMFA). Violin plots show areas under the LMFA and GMFA curves (AUC) between 80 and 200 ms. Statistical differences between conditions are denoted with asterisks (*n.s*.: not significant). Topographic plots of EEG responses from the contralateral motor cortices, corresponding to panels DEHI, are shown in Fig. S4 AB. Signals are averaged over 23 (ACG), 21 (DH), 14 (EI), and 56 (BFJ) epochs and shown along 95% confidence intervals (C to J).

## 3. Results

### 3.1. Real-time IED-detection performance

Of TMS pulses aimed at an IED in all stimulation locations, 19% (P1) and 39% (P2) were true positives, *i.e*., successfully delivered during an IED (P1: 87/465, 95%-CI 15−23%, P2: 137/347, CI 34−45%). Of pulses aimed at non-IEDs, 99.8% (P1) and 99.7% (P2) were true negatives, *i.e*., successfully delivered during a non-IED period (P1: 459/460, CI 99−100%, P2: 345/346, CI 98−100%). The probability of achieving equal or better IED-targeting success rate with randomly timed TMS is < 10^−30^ (Supplementary Material).

### 3.2. TEP results

P1 had predominantly left-hemisphere IEDs, and P2 had right-hemisphere IEDs. We present TEP results based on whether stimulation was ipsilateral (left hemisphere for P1, right for P2) or contralateral (right hemisphere for P1, left for P2) to IEDs.

In P1, AUC was larger in the IED−TEP condition than the nTEP or IED-alone conditions for ipsilateral M1 (LMFA and GMFA; Fig. 2 ACG) and ipsilateral occipital stimulation (LMFA only; Fig. S3 C), suggesting the IED modified the TMS response at these sites. AUC was smaller in the nTEP condition than the IED−TEP or IED-alone condition for contralateral M1 (LMFA and GMFA: Fig. 2 DH), contralateral occipital (GFMA only; Fig. S3 G), and ipsilateral occipital (GMFA only; Fig. S3 H) stimulation, suggesting the IED itself drives AUC differences at these sites. Statistics are in Supplementary Tables S2 and S3.

In P2, AUC was smaller in the nTEP condition than either the IED−TEP or IED-alone conditions for ipsilateral M1 (LMFA only; Fig. 2 F) and both ipsilateral and contralateral occipital stimulation (GMFA only; Fig. S3 IJ), again suggesting AUC differences were driven by IED itself at these sites. There were no other differences in AUC.

## 4. Discussion

We developed a real-time IED-detection algorithm and timed TMS to IEDs across four stimulation locations in two participants. We developed a preprocessing pipeline for TMS−EEG data with IEDs and found that IED-timed TMS yields physiological insights.

### 4.1. Development of the IED-detection algorithm

The IED-detection algorithm succeeded in triggering TMS before the end of an IED, a task requiring rapid signal processing. The consistency of TMS delivery related to the IED morphology was high. Although the IED-targeting success rates were relatively low (19−39%), such rates could not be achieved with randomly given TMS. The algorithm performance can be improved further by increasing training dataset size, making it more robust to variability of IED morphology within participants.

### 4.2. Effects of IED-timed TMS on TEPs

IEDs modulated late TEPs in one participant in a site-specific manner. In P1, late TEPs were much larger when TMS was administered concurrently with an IED than during a non-IED period. Notably, this was observed when stimulating the hemisphere generating IEDs, but not when stimulating the contralateral hemisphere. In contrast, IEDs did not modulate TEPs for P2.

Several explanations exist for differences in participant responses. Stimulation intensity likely plays an important role, as we stimulated with suprathreshold intensity only in P1. Suprathreshold stimulation may be necessary to sufficiently perturb the cortical network during an IED. Suprathreshold stimulation also induces larger sensory-evoked potentials, which can contribute to the late TEP.

Antiseizure medications may additionally impact results. P2 took a voltage-gated sodium channel blocker (oxcarbazepine), medication that stabilizes membrane potential, increases rMT, and changes TEP amplitudes (Darmani et al., 2019). Chronic membrane potential stabilization from sodium channel blockade may dampen both cortical response to TMS and transient hyperexcitability during IEDs, reducing dynamic range between IED and non-IED states and resulting in attenuated state-dependent TEP modulation.

Understanding the mechanistic overlap between IED slow-waves and TMS-evoked N100s is critical as both occur within the same temporal window (80−200 ms) and may reflect related inhibitory processes. The post-IED slow-wave represents a refractory period with high inhibition (Dorn & Witte, 1995), while the TMS-evoked N100 reflects a balance between gamma-aminobutyric acid (GABA)-inhibitory and glutamate-excitatory levels (Du et al., 2018). When delivered during an IED, exogenous TMS-evoked inhibition interacts with the endogenous IED-evoked inhibitory state. The resulting late TEP amplitude may reflect the capacity of the epileptic network to recruit additional inhibitory resources when perturbed during an IED. A key question in epilepsy is whether IEDs are pro- or anti-convulsant, and IED-timed TMS may be an important tool to study this open question.

IED-timed TMS offers the opportunity to study epilepsy networks non-invasively. IED propagation involves sequential activation of cortical nodes, where early (“upstream”) nodes initiate discharges and late (“downstream”) nodes represent terminal propagation sites. Invasive electrical stimulation of upstream nodes elicits broad network responses, while stimulation of intermediate or downstream nodes produces attenuated responses (Tomlinson et al., 2025). For P1, left M1 likely represents an upstream node. When we delivered suprathreshold TMS to left M1 during an IED, we observed an amplified late TEP that was both site-specific and globally evident. This supports prior evidence that the epileptic network exists in a hypersynchronous, hyperconnected configuration during the IED (Goad et al., 2022), priming the network for exaggerated responses to perturbation. When suprathreshold TMS engages this primed upstream node, it may trigger a widespread compensatory inhibitory surge, manifested as an enlarged N100. This could represent a “safety mechanism” wherein dual perturbation (endogenous IED + exogenous TMS) recruits strong inhibitory circuits to prevent escalation toward seizure, consistent with the role of the IED slow-wave in limiting seizure propagation (Dorn & Witte, 1995). In contrast, P2 had IEDs with a more temporal distribution, suggesting that M1 was a downstream node. IED−TEPs were not distinguishable from IEDs alone, consistent with attenuated responses reported during retrograde stimulation from downstream nodes (Tomlinson et al., 2025). Delivering TMS to downstream nodes may not recruit additional network-wide responses because spontaneous IEDs have already engaged relevant pathways. IED-timed TMS may thus be a valuable tool for deciding upon best cortical targets for therapeutic neurostimulation.

### 4.3. Future directions

Several approaches could improve IED-targeting success rate and consistency. Adjusting the sensitivity–specificity trade-off through a three-class system (“IED,” “non-IED,” “uncertain”) could reduce false positives by triggering TMS only when the classifier is highly certain. Additional training data would improve generalization, particularly if diverse IED morphologies across arousal states are included. Real-time low-latency filtering could reduce noise without introducing detection delays. To improve temporal precision, the detection algorithm could focus on the initial rising phase of the IED, enabling more consistent stimulation timing. Alternatively, simpler amplitude-threshold detection methods may work for syndromes with high-amplitude IEDs. More sophisticated methods could potentially anticipate IED onset, although validating predictions would be challenging if TMS alters upcoming IEDs.

In future research, studies of larger cohorts with inclusion of structural brain connectivity data will be needed to characterize individual variability in state-dependent responses. Methodological improvements should include target optimization (Ukharova et al., 2025) to minimize TMS-induced artifacts during measurements, and assessment of multiple TMS intensities to clarify the threshold for state-dependent modulation. In data processing, deduction or signal-space projection (Mutanen et al., 2022) could separate IED and TEP components for clearer interpretation of their interaction.

IED-timed TMS has potential applications in both clinical care and epilepsy research. For presurgical evaluation, the method could map epileptogenic networks non-invasively, complementing current invasive mapping approaches, while therapeutically brain-state-dependent rTMS targeting could reduce treatment variability and improve efficacy (VanHaerents et al., 2020). The presented method should be tested in medically refractory epilepsy syndromes where new treatment options are urgently needed (Specchio et al., 2022). Furthermore, network responses during IED-timed TMS offers a unique opportunity to address fundamental questions about whether IEDs have anti-convulsant properties and how IEDs impact brain networks.

## 5. Conclusion

Timing TMS to IEDs is feasible. IEDs modulate the late TEP in a participant- and site-specific manner. Increasing the IED-targeting success rate of the real-time algorithm as well as performing measurements in more participants are critical next steps. The presented method shows promise as a non-invasive protocol for investigating transient epileptic brain states and optimizing neurostimulation therapies.

## CRediT authorship contribution statement

**Matilda Makkonen:** Conceptualization, Data curation, Formal analysis, Funding acquisition, Investigation, Methodology, Visualization, Writing – original draft, Writing – review and editing. **Olli-Pekka Kahilakoski:** Resources, Software, Writing – review and editing. **Miguel Menchaca:** Data curation, Investigation, Writing – review and editing. **Ivan Zubarev:** Methodology, Resources, Software, Writing – review and editing. **Oula Siljamo:** Software, Writing – review and editing. **Umair Hassan:** Resources, Writing – review and editing. **Xiwei She:** Data curation, Investigation, Writing – review and editing. **Wendy Qi:** Data curation, Investigation, Writing – review and editing. **Tuomas P. Mutanen:** Funding acquisition, Supervision, Writing – review and editing. **Risto J. Ilmoniemi:** Funding acquisition, Supervision, Writing – review and editing. **Pantelis Lioumis:** Conceptualization, Funding acquisition, Methodology, Project administration, Supervision, Writing – review and editing. **Fiona M. Baumer:** Conceptualization, Data curation, Funding acquisition, Investigation, Methodology, Project administration, Supervision, Writing – original draft, Writing – review and editing.

## Data availability

The full data is available from the corresponding author on reasonable request.

## Declaration of generative AI and AI-assisted technologies in the writing process

During the preparation of this work the authors used ChatGPT-4o and Anthropic Claude Sonnet 4.5 in order to edit and proofread the language of the raw text. After using this tool, the authors reviewed and edited the content as needed and take full responsibility for the content of the published article.

## Acknowledgments

We are grateful for the participants and their families, without whom this study would not have been possible. This study was enabled with funding from Emil Aaltonen Foundation (MM; grant number 240110 K1), Aalto University US Global Program Pilot, and Finnish Cultural Foundation (MM; grant number 00250619), and has also been supported by the European Research Council (ERC Synergy) under the European Union’s Horizon 2020 research and innovation programme (ConnectToBrain; grant agreement No 810377), the K23 Career Development Award (NINDS K23NS116110) (FMB), and the Wu Tsai Neuroscience Institute Koret Human Neuroscience Community Lab Pilot Award. We thank Olivia Peony and Miles Lucas for support with handling data, Lotta Makkonen for the illustration in the graphical abstract, as well as Ilkka Rissanen and Elena Ukharova for discussing important concepts.

