## Supplementary Material for "TMS timed to interictal epileptiform discharges"

[a]*Department of Neuroscience and Biomedical Engineering, Aalto University, Espoo, Finland*
[b]*Department of Neurology, Stanford University School of Medicine, Stanford, CA, USA*
[c]*Department of Psychiatry, Stanford University School of Medicine, Stanford, CA, USA*
[d]*BioMag Laboratory, HUS Medical Imaging Center, Aalto University, University of Helsinki and Helsinki University Hospital, Helsinki, Finland*
[e]*Cognitive Brain Research Unit, Department of Psychology and Logopedics, Faculty of Medicine, University of Helsinki, Helsinki, Finland*
[*]*Shared last authorship*


### S1. Interictal epileptic discharge (IED)-detection algorithm: training and implementation

This section provides detailed information about the machine learning algorithm referenced in Main Text Section 2.2.

#### S1.1. Training data acquisition

Sessions during which training data was acquired were conducted separately from the experimental transcranial magnetic stimulation (TMS) sessions. For P1, four resting-state electroencephalography (EEG) sessions (totaling 1134 s) were recorded on two consecutive days. For P2, three EEG sessions (totaling 959 s) were recorded on a single day. Some TMS pulses were administered during P2's training session, but all data surrounding the pulses were removed from further processing, as explained later. The TMS pulse artifacts were also removed before downsampling to avoid filtering artifacts: the signal was zeroed from 2 ms before to 10 ms after the pulse onset and replaced with a linear interpolation of the surrounding signal.

#### S1.2. IED annotation from training data

A pediatric epileptologist (FMB) annotated IED onset times using the built-in annotation tool of the MNE-Python software (Gramfort et al., 2013). Data were displayed in a clinically standard bipolar montage to optimize IED visibility. No other preprocessing was applied before annotation. In total, 327 IEDs (P1) and 498 IEDs (P2) were annotated. The signal from 10 ms before to 120 ms after each

IED annotation was marked as IED. IEDs thus represented 3.7% (P1) and 6.8% (P2) of the total training data duration.

#### S1.3. Segment creation and labeling

Following downsampling to 1 kHz and conversion to common average reference, the continuous data were divided into 130-sample (130-ms) segments with 50% overlap to increase the number of training examples. This segment length was chosen to capture complete IEDs. Each segment was baseline-corrected by removing the segment mean in each channel and scaled by dividing each value in the segment by the standard deviation over all channels and time points. For P2, the segments including at least one sample from 5 ms before to 200 ms after a TMS pulse were discarded to make the data as similar to resting-state EEG as possible.

IED detection was defined as a binary classification task. A segment was labeled "IED" if > 45% of its duration overlapped with an IED annotation, and "non-IED" otherwise. In total, P1 had 17469 segments, of which 663 (3.8%) were labeled "IED." P2 had 14764 segments, of which 1542 (10.5%) were discarded due to TMS pulses, and 1033 (7.8%) of the remaining segments were labeled "IED." Class distribution was thus highly imbalanced for both participants.

#### S1.4. Class balancing

Class imbalance can impair neural network training. We used rejection resampling, in which non-IED segments were randomly discarded until both classes had equal representation in each training batch. This approach was chosen over synthetic minority oversampling to avoid overfitting. The data were divided into 5 folds, one of which was chosen as a validation set. The validation set maintained natural class proportions to provide realistic performance estimates.

#### S1.5. Linear-filter CNN architecture and training

The LF-CNN model (Zubarev et al., 2019) was selected for IED detection as it is designed for classifying magnetoencephalography (MEG) and EEG data. The model has only a few layers and few trainable parameters compared to many other CNN classifiers, reducing the risk of overfitting to the noisy M/EEG data and decreasing the amount of training data required.

The main architecture parameters were as follows:

- Number of latent sources: 32 (recommended by Zubarev et al., 2019)
- Regularization: $L^1$ on filter weights, dropout on output layer ($L^2$ regularization not applied)
- Padding: "Same" padding on the convolutional layer input
- Output nonlinearity: Softmax for binary classification
- Optimizer: Adam (Kingma & Ba, 2014), minimizing multinomial cross-entropy

During parameter search, the LF-CNN model was trained for each participant separately for a single fold with a class-balanced test set of 100 EEG segments. The performance was evaluated on the validation set after every iteration. Early stopping prevented overfitting: training terminated when validation loss plateaued for 5 (P1) or 3 (P2) consecutive iterations; the model for P2 started to overfit faster.

Model hyperparameters were tuned separately for both participants, with validation accuracy as the target variable. Initial values were chosen based on Makkonen (2023), which presented real-time IED detection from EEG of self-limited epilepsy with centrotemporal spikes (SeLECTS) patients. This detection was developed on data from a different patient than the participants in the present study. The hyperparameters leading to the best validation accuracy are shown in Table S1. The model training and parameter optimization were performed with the MNEflow package (Zubarev et al., 2022).

**Table S1:** The optimal hyperparameter values for IED-detecting machine-learning model LF-CNN training.

| **Parameter** | **Value** |
|---|---|
| Temporal (convolutional) filter length | 128 |
| Learning rate | 0.005 (P1), 0.05 (P2) |
| $L^1$ regularization scalar | 0.001 (P1), 0.01 (P2) |
| Pooling factor (max-pooling) | 64 |
| Pooling stride | 64 |
| Dropout coefficient | 0.1 |
| Hidden layer activation function | Rectified linear unit (ReLU) |
| Train-batch size | 100 |

### S1.6. Offline performance testing

The performance of the trained models was tested on EEG datasets that were not included in the training data but were collected in the same measurement session(s). For P1, the testing data consisted of three resting-state EEG recordings with a total duration of 620 s and 74 annotated IEDs. For P2, the testing data consisted of five TMS–EEG recordings with a total duration of 1586 s and 646 IEDs. As with the training data, TMS pulse artifacts were removed: the signal was zeroed from 2 ms before to 10 ms after the pulse onset and replaced with a linear interpolation of the surrounding signal.

Model testing was performed in a mock-real-time scenario, in which the continuous EEG signal was divided into 130-ms segments with 110-ms overlap, *i.e.*, a 20-ms shift between segments. As with the training data, each segment was scaled to baseline. If the model classified a segment as an IED, the segment end time was stored as an IED-event and used as a proxy for pulse timing in the real-time experiment, in which a pulse was given only after receiving the classification result of a segment.

The detection performance was assessed by comparing the IED-event timestamps to the start times of the annotated IEDs. True IED detections were defined as an IED-event occurring no longer than 230 ms after the start of an IED. False-positive detections were defined as an IED-event occurring at any other time. The 230-ms threshold was chosen to account for the IED duration and segment length. When the distance of two consecutive IED-events was less than 60 ms, only the first event of this "burst" was included in the analysis, simulating the inter-stimulus interval of the real-time experiment. False negatives were defined as IEDs without an IED-event within the 230-ms time window. For each testing data set, we calculated the proportion of false-positive IED detections out of all IED detections, and the false-negative rate, *i.e.*, the proportion of false-negative IED detections (missed IEDs) out of all annotated IEDs.

For P1, the best-performing model yielded 36% false-positive IED detections and a false-negative rate of 5.8%, *i.e.*, missed 5.8% of the IEDs. For P2, the best-performing model gave 32% false-positive IED-perceptions and a false-negative rate of 1.2%. These metrics guided the model selection for real-time deployment. The detection latencies (median, minimum, and maximum) were 62, 13, and 229 ms (P1), and 59, 21, and 138 ms (P2).

#### S1.7. Real-time integration

During the experiment, a TurboLink device (Brain Products GmbH, Germany) streamed EEG in real time from the amplifier, one sample at a time, to a laptop computer (Dell Precision 7680) with a 24-core Intel Core i9-13950HX processor, 32 GB of RAM and the Ubuntu 22.04 LTS operating system with the PREEMPT_RT real-time kernel patch.

The laptop computer was running the NeuroSimo software (Kahilakoski et al., 2025), which maintained a buffer of the latest 130 samples (130 ms) and classified the latest 130-sample segment every 20 ms with the pre-trained LF-CNN model. Before classification, each segment was baseline-corrected by removing the mean of the segment in each channel and scaled by dividing each value in the segment by the standard deviation over all channels and time points. Based on the classification, a TMS pulse was delivered by generating and transmitting a trigger signal to the TMS device via a LabJack T4 USB peripheral.

### S2. Experimental setup: additional details

This section expands on Main Text Section 2.3. The minimum ISIs were chosen to balance between obtaining reasonably segregated states and obtaining an adequate number of epochs of each state in a reasonable time; this was a concern particularly for the IED stimuli as we could not pre-specify the number of IEDs a participant would have. The minimum ISI for P1 was 2 s. Before measuring P2, we refined the ISI approach based on our experience of the P1 measurement. We reduced the minimum ISI to 1 s before IED-targeting pulses to shorten the measurement. In addition, non-IED-targeting pulses were delivered only when at least 1.5 s had passed from the previous real-time IED detection. This ensured that non-IED epochs were not contaminated by recent IED activity.

During the experiment, the participants were seated in a comfortable semi-reclined position. The electrodes were prepared with conductive gel, and their impedances were monitored and kept below 5 kΩ. To mask the auditory brain response to TMS pulses, the participants wore in-ear headphones playing noise matched in frequency to the TMS clicks. The volume of the noise was set to a safe level so that the participants could not hear the TMS coil clicks. Over-ear headphones were placed on top to further muffle noise from the room.

For P1, a total of 925 pulses were administered, with four 99-pulse blocks to the left primary motor cortex (M1) and two blocks to the right M1, right occipital cortex and left occipital cortex each, in this order. The fourth left-M1 stimulation block contained only 34 pulses as the block had to be stopped early due to the participant needing to change the seating position. For P2, 693 pulses were administered in seven blocks in the following order: RM, RM, LM, RM, LM, RO, LO, where R and L indicate right and left hemisphere and M and O indicate M1 and occipital stimulation site, respectively. The first left-M1 block was discarded from TEP analysis due to the participant falling asleep during the block. Algorithm performance data from this block were retained as they did not depend on wakefulness.

### S3. Preprocessing details

Preprocessing was performed with the TESA plugin (Rogasch et al., 2017) of EEGLAB (Delorme and Makeig, 2004) in Matlab 2022a (The Mathworks, Inc., MA, USA). Robust detrending (de Cheveigné and Arzounian, 2018) was performed with a second-order polynomial function.

In independent component analysis (ICA), the data were compressed to 30 dimensions with principal component analysis (PCA) and then decomposed into 30 independent components with FastICA (Hyvärinen and Oja, 1997). The components corresponding to eye blinks and horizontal eye movement were manually identified based on topography and waveform and removed from the data. Between one and three such components were identified and removed from each data set. The data were baseline corrected after ICA, setting 500 to 70 ms pre-stimulus as the baseline.

The regularization parameter of the source-estimate-utilizing noise-discarding algorithm (SOUND; Mutanen et al., 2022) was set to 0.01, as larger values of the parameter led to IED amplitude suppression. The custom lead-field matrix for SOUND was computed using average electrode locations. Signal-space-projection–source-informed reconstruction (SSP–SIR; Mutanen et al., 2022) was used to remove four principal components. The data were baseline-corrected and re-referenced to global average at the end of the pipeline.

M1 ROIs were six-electrode groups including the stimulation site and the most prominent IED electrode, while occipital ROIs comprised four-electrode groups closest to the stimulation site.

### S4. Epoch classification diagrams and performance calculation

Figures S1 and S2 provide a visual flowchart of epoch selection and classification procedures described in Main Text Sections 2.4.1. and 2.4.2.1.

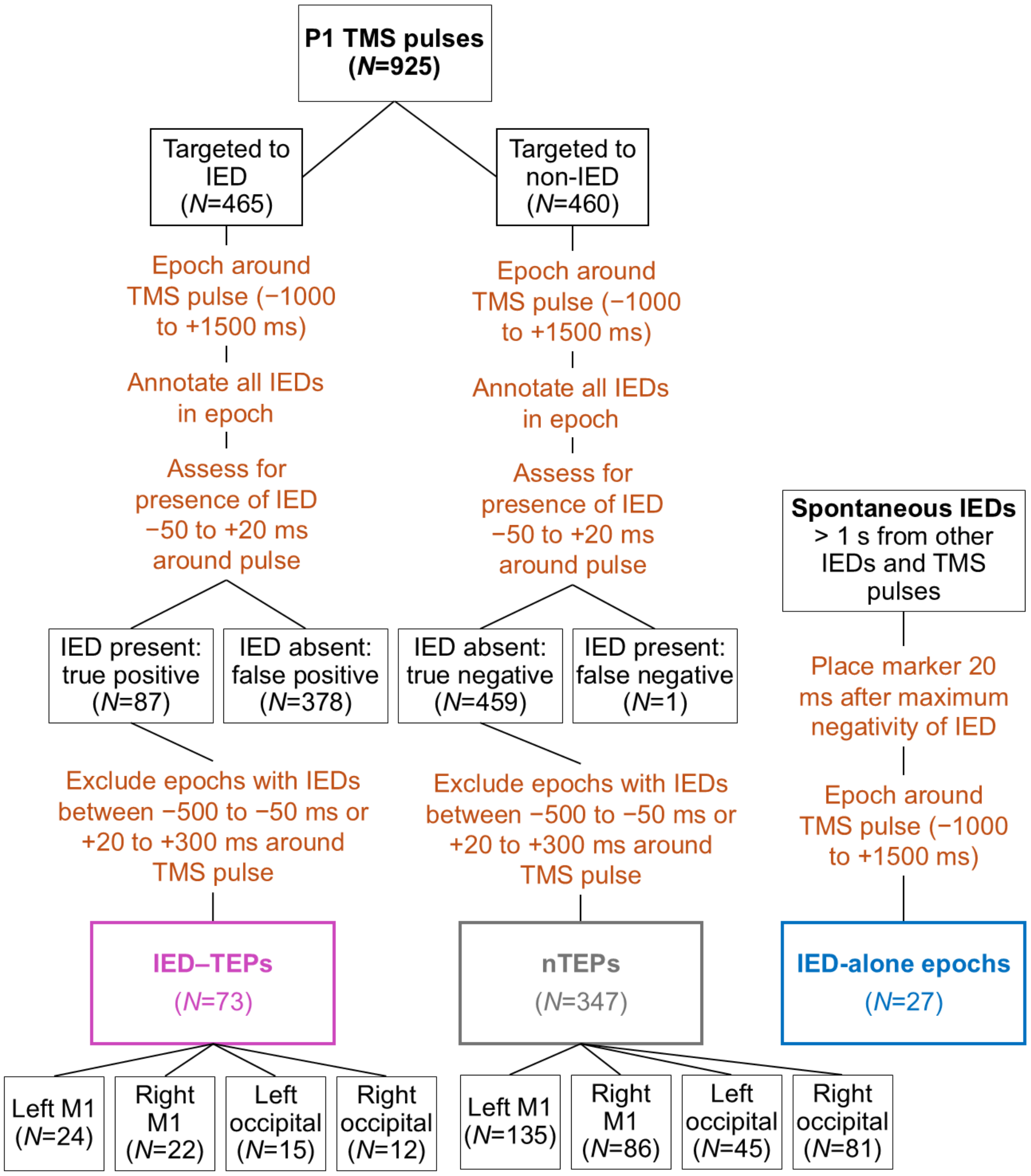


**Figure S1:** Diagram presenting epoch processing and selection before preprocessing for P1. The IED-alone data were extracted from the left M1 dataset.

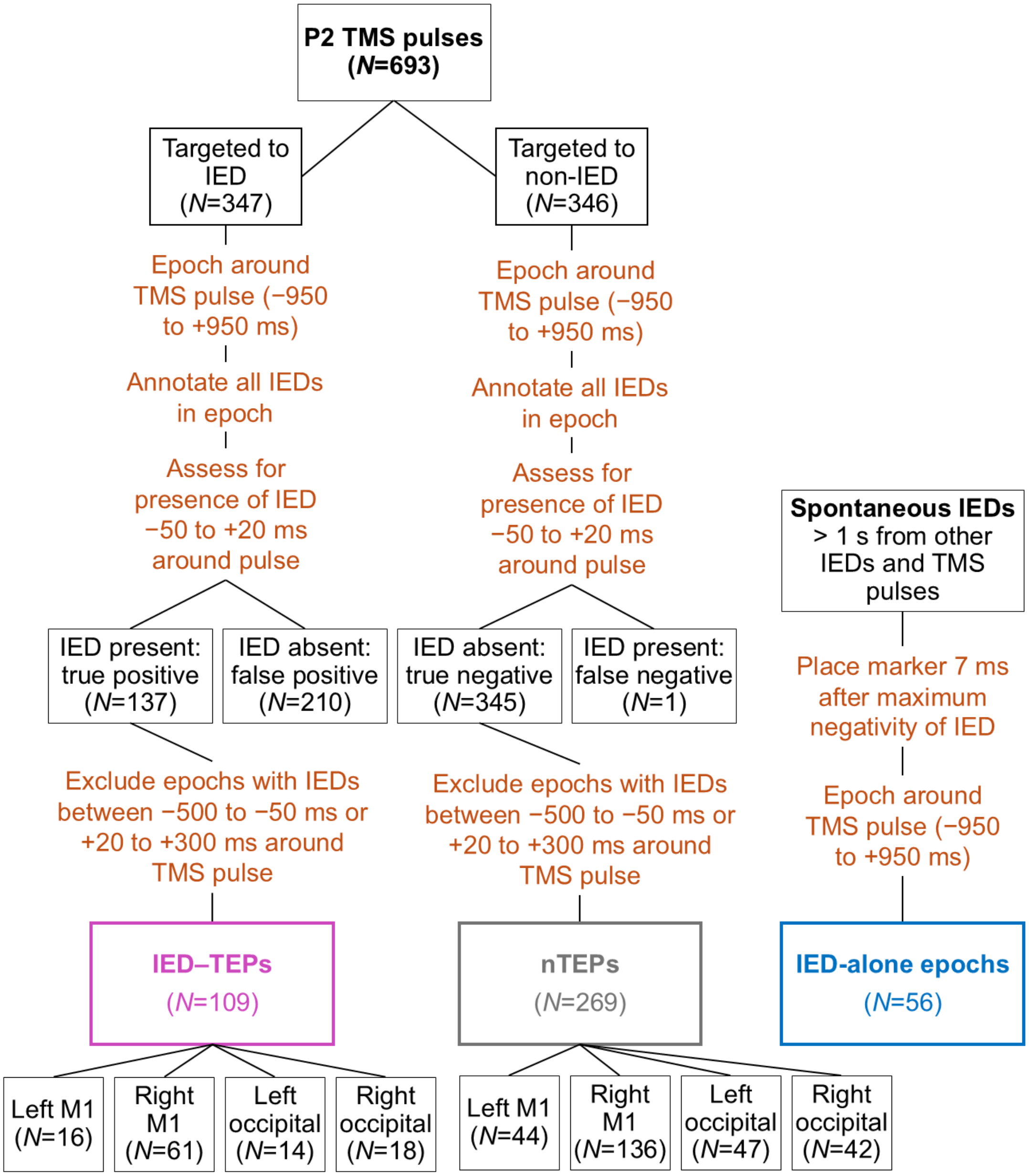


**Figure S2:** Diagram presenting epoch processing and selection before preprocessing for P2. The IED-alone data were extracted from the right M1 dataset.

#### S4.1. Statistical comparison to random stimulation

To evaluate whether the algorithm performed better than randomly timed TMS, we calculated the theoretical probability $P_r$ for achieving an IED-targeting success rate equal to or greater than in our results with randomly given TMS (pulse times sampled from a uniform distribution over the session duration). This can be calculated with the binomial probability of repeated events:

$$P_r = \sum_{k=n_a}^{N} \binom{N}{k} (p_r)^k (1 - p_r)^{N-k}, \text{(S1)}$$

where $N$ is the total number of pulses targeted at an IED, $n_a$ is the number of pulses that successfully hit an IED, and $p_r$ is the probability of one randomly timed pulse to hit an IED, *i.e.*, the fraction of the EEG data marked as IEDs, calculated from the training data.

For P1: $N$ = 465, $n_a$ = 87, $p_r$ = 0.037 → $P_r = 2.2 \cdot 10^{-35}$

For P2: $N$ = 347, $n_a$ = 137, $p_r$ = 0.068 → $P_r = 2.6 \cdot 10^{-67}$

This means that achieving our observed IED-hit rates by randomly timing TMS pulses is essentially impossible with the same number of TMS pulses as in our measurements.

### S5. Supplementary results

Tables S2 and S3 present the statistics for AUC comparisons between conditions for P1 and P2, respectively. Fig. S3 shows the TEPs, LMFA, and GMFA from the occipital stimulation sites. Fig. S4 shows the channel-wise TEPs for IED-contralateral M1 and occipital stimulation, corresponding to panels D, E, H, and I in Figs. 2 and S3.

**Table S2:** Statistics from AUC comparisons between conditions from P1. If the Kruskal-Wallis (KW) non-parametric test returned significant results ($p$ < 0.05, indicated with yellow), the Dunn–Šidák post-hoc test was performed. If the KW test was not significant ($p$ > 0.05), the post-hoc test was not performed (n.p.). *Contra* and *ipsi* indicate whether stimulation was ipsilateral (left hemisphere) or contralateral (right hemisphere) to IEDs.

| Stimulation location | Metric | Condition | AUC median & inter-quartile range | Kruskal-Wallis (KW) $\chi 2$ | KW $p$ value | Post-hoc $p$ vs. nTEP | Post-hoc $p$ vs. IED |
|---|---|---|---|---|---|---|---|
| Right (contra) motor | LMFA | IED–TEP | 1786 [1008–2285] | $\chi 2(2,60)$= 8.9 | 0.011 | 0.032 | 1.0 |
| Right (contra) motor | LMFA | nTEP | 1079 [850–1305] | | | | 0.026 |
| Right (contra) motor | LMFA | IED | 1709 [963–2557] | | | | |
| Right (contra) motor | GMFA | IED–TEP | 1930 [1251–2821] | $\chi 2(2,60)$= 24 | 6.1E−6 | 6.6E−5 | 1.0 |
| Right (contra) motor | GMFA | nTEP | 1002 [837–1189] | | | | 6.6E−5 |
| Right (contra) motor | GMFA | IED | 1836 [1469–2901] | | | | |
| Left (ipsi) motor | LMFA | IED–TEP | 5686 [3591–6741] | $\chi 2(2,66)$= 24 | 5.2E−6 | 1.5E−5 | 2.9E−4 |
| Left (ipsi) motor | LMFA | nTEP | 1537 [1411–2040] | | | | 0.87 |
| Left (ipsi) motor | LMFA | IED | 2072 [1029–2895] | | | | |
| Left (ipsi) motor | GMFA | IED–TEP | 3540 [2415–4332] | $\chi 2(2,66)$= 31 | 1.6E−7 | 8.6E−8 | 0.0022 |
| Left (ipsi) motor | GMFA | nTEP | 1398 [1118–1542] | | | | 0.088 |
| Left (ipsi) motor | GMFA | IED | 1846 [1316–2315] | | | | |

| | | | | | | | |
|---|---|---|---|---|---|---|---|
| Right (contra) occipital | LMFA | IED–TEP | 1152 [1024–1442] | $\chi 2(2,33)=$ 4.4 | 0.11 | n.p. | n.p. |
| Right (contra) occipital | LMFA | nTEP | 1004 [873–1076] | | | | n.p. |
| Right (contra) occipital | LMFA | IED | 1084 [834–1220] | | | | |
| Right (contra) occipital | GMFA | IED–TEP | 2138 [1423–3168] | $\chi 2(2,33)=$ 17 | 2.2E–4 | 7.0E–4 | 0.99 |
| Right (contra) occipital | GMFA | nTEP | 904 [860–985] | | | | 0.0019 |
| Right (contra) occipital | GMFA | IED | 2022 [1113–3271] | | | | |
| Left (ipsi) occipital | LMFA | IED–TEP | 1631 [1327–2183] | $\chi 2(2,33)=$ 9.8 | 0.0073 | 0.012 | 0.033 |
| Left (ipsi) occipital | LMFA | nTEP | 1156 [964–1386] | | | | 0.98 |
| Left (ipsi) occipital | LMFA | IED | 1188 [979–1465] | | | | |
| Left (ipsi) occipital | GMFA | IED–TEP | 2031 [1458–2936] | $\chi 2(2,33)=$ 17 | 1.6E–4 | 0.0018 | 0.98 |
| Left (ipsi) occipital | GMFA | nTEP | 893 [826–1315] | | | | 4.7E–4 |
| Left (ipsi) occipital | GMFA | IED | 2051 [1657–3087] | | | | |

**Table S3:** Statistics from AUC comparisons between conditions from P2. If the Kruskal-Wallis (KW) non-parametric test returned significant results ($p < 0.05$, indicated with yellow), the Dunn–Šidák post-hoc test was performed. If the KW test was not significant ($p > 0.05$), the post-hoc test was not performed (n.p.). *Contra* and *ipsi* indicate whether stimulation was ipsilateral (right hemisphere) or contralateral (left hemisphere) to IEDs.

| **Stimulation location** | **Metric** | **Condition** | **AUC median & inter-quartile range** | **Kruskal-Wallis (KW) $\chi 2$** | **KW *p* value** | **Post-hoc *p* vs. nTEP** | **Post-hoc *p* vs. IED** |
|---|---|---|---|---|---|---|---|
| Right (ipsi) motor | LMFA | IED–TEP | 1789 [1283–2269] | $\chi 2(2,165)=$ 12 | 0.0029 | 0.11 | 0.48 |
| Right (ipsi) motor | LMFA | nTEP | 1462 [1187–1787] | | | | 0.0021 |
| Right (ipsi) motor | LMFA | IED | 1882 [1583–2343] | | | | |

| | | | | | | | |
|---|---|---|---|---|---|---|---|
| Right (ipsi) motor | GMFA | IED–TEP | 1763 [1455–2175] | $\chi2(2,165)=3.0$ | 0.23 | n.p. | n.p. |
| Right (ipsi) motor | GMFA | nTEP | 1669 [1358–2071] | | | | n.p. |
| Right (ipsi) motor | GMFA | IED | 1782 [1603–2035] | | | | |
| Left (contra) motor | LMFA | IED–TEP | 1157 [855–1477] | $\chi2(2,39)=4.1$ | 0.13 | n.p. | n.p. |
| Left (contra) motor | LMFA | nTEP | 1498 [1208–2021] | | | | n.p. |
| Left (contra) motor | LMFA | IED | 1085 [944–1492] | | | | |
| Left (contra) motor | GMFA | IED–TEP | 1794 [1313–2285] | $\chi2(2,39)=0.79$ | 0.67 | n.p. | n.p. |
| Left (contra) motor | GMFA | nTEP | 1579 [1277–1959] | | | | n.p. |
| Left (contra) motor | GMFA | IED | 1820 [1694–1976] | | | | |
| Right (ipsi) occipital | LMFA | IED–TEP | 1282 [1122–1731] | $\chi2(2,51)=5.1$ | 0.079 | n.p. | n.p. |
| Right (ipsi) occipital | LMFA | nTEP | 1177 [973–1680] | | | | n.p. |
| Right (ipsi) occipital | LMFA | IED | 1652 [1312–1870] | | | | |
| Right (ipsi) occipital | GMFA | IED–TEP | 1600 [1287–2064] | $\chi2(2,51)=6.3$ | 0.044 | 0.33 | 0.71 |
| Right (ipsi) occipital | GMFA | nTEP | 1507 [1081–1783] | | | | 0.039 |
| Right (ipsi) occipital | GMFA | IED | 1769 [1629–2007] | | | | |
| Left (contra) occipital | LMFA | IED–TEP | 1524 [1046–2683] | $\chi2(2,33)=0.43$ | 0.81 | n.p. | n.p. |
| Left (contra) occipital | LMFA | nTEP | 1539 [1292–1878] | | | | n.p. |
| Left (contra) occipital | LMFA | IED | 1703 [1415–2343] | | | | |
| Left (contra) occipital | GMFA | IED–TEP | 1670 [1530–2617] | $\chi2(2,33)=11$ | 0.0038 | 0.083 | 0.62 |
| Left (contra) occipital | GMFA | nTEP | 1433 [1274–1674] | | | | 0.0032 |
| Left (contra) occipital | GMFA | IED | 1960 [1718–2525] | | | | |

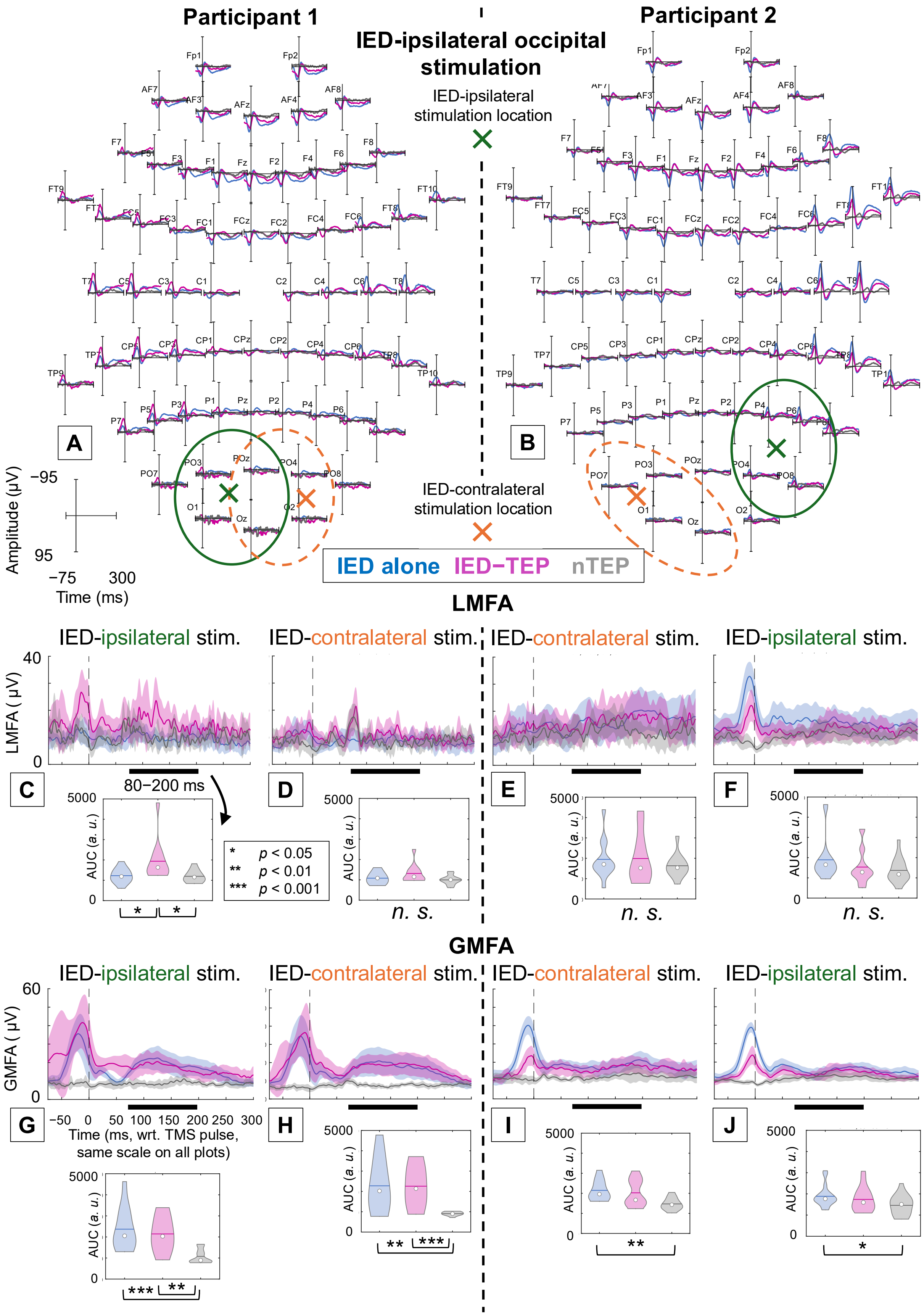

Participant 1
IED-ipsilateral occipital stimulation
Participant 2
IED-ipsilateral stimulation location
IED-contralateral stimulation location
A
B
Amplitude (μV)
−95
95
−75
300
Time (ms)
IED alone
IED−TEP
nTEP
LMFA
IED-ipsilateral stim.
IED-contralateral stim.
IED-contralateral stim.
IED-ipsilateral stim.
LMFA ( μV)
40
0
80−200 ms
C
D
E
F
AUC (a. u.)
5000
0
* p < 0.05
** p < 0.01
*** p < 0.001
n. s.
n. s.
n. s.
GMFA
IED-ipsilateral stim.
IED-contralateral stim.
IED-contralateral stim.
IED-ipsilateral stim.
GMFA ( μV)
60
0
G
H
I
J
-50 0 50 100 150 200 250 300
Time (ms, wrt. TMS pulse, same scale on all plots)

**Figure S3:** Panels A and B: Topographic plots of EEG signals in each channel in the global average reference for IEDs (blue), and TMS-evoked responses when successfully targeting an IED (pink) and a non-IED (nTEP, grey) for P1 (A) and P2 (B); stimulation applied to occipital cortex in the IED-generating (ipsilateral) hemisphere (A: left, B: right). Stimulation sites are presented as crosses with ROI electrodes circled. Green cross and circle represent ipsilateral stimulation and ROI, respectively, while orange cross and circle represent the contralateral counterparts. Panels C to F: Local mean field amplitudes (LMFA) of the four-electrode ROI. Panels G to J: Global mean field amplitudes (GMFA). Violin plots show areas under the LMFA and GMFA curves (AUC) between 80 and 200 ms. Statistical differences between conditions are denoted with asterisks (*n.s.*: not significant). Topographic plots of EEG responses from the IED-contralateral occipital cortices, corresponding to panels DEHI, are shown in Fig. S4 CD. The signals are averaged over 12 (ACDEGHI), and 18 (BFJ) epochs and shown along 95% confidence intervals (C to J).

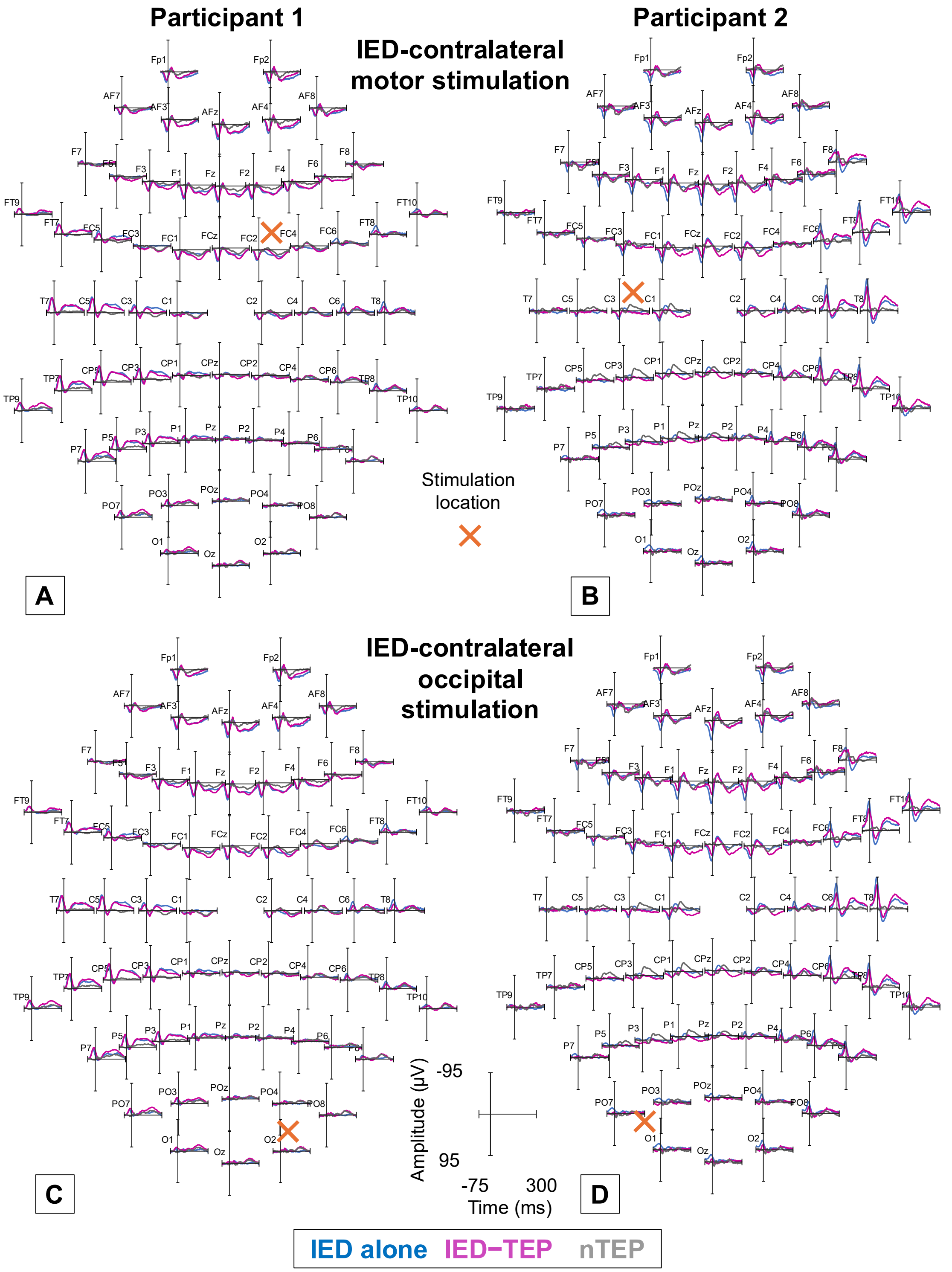


**Figure S4:** Topographic plots of signals in each EEG channel in the global average reference for IEDs (blue), and TMS-evoked responses when targeting an IED (pink) or a non-IED (nTEP; grey) for participants 1 (AC) and 2 (BD) when stimulating the contralateral (P1: right, P2: left) motor (AB) and occipital (CD) cortex from the IED-generating hemisphere. The signals are averaged over 21 (A), 14 (B), and 12 (CD) epochs.
